# Multi-omic analysis of vaginal bacteria:host immune and metabolic interactions in adverse early pregnancy outcomes

**DOI:** 10.1101/2025.08.11.669732

**Authors:** Shabnam Bobdiwala, Pamela Pruski, Gonçalo DS Correia, Maya Al-Memar, Yun S. Lee, Ann Smith, Julian R. Marchesi, Phillip R. Bennett, Zoltán Takáts, Tom Bourne, David A. MacIntyre

**Author notes:** S.B, P.P. and G.DS.C. contributed equally to this study. The order of authorship was determined based on the chronological progression of the investigation. S. B. performed clinical sample collection and the metataxonomic experiments, P.P. mass spectrometry metabolomics, metabolite identification, and immune profiling experiments, and G.C. bioinformatic analysis. All co-first authors contributed equally to writing of the manuscript. corresponding author: David A. MacIntyre. Disclosure Statement: P.P., Z.T., P.R.B and D.A.M hold patents for the use of rapid evaporative ionization mass spectrometry and DESI-MS analysis of swabs and biopsies (US10026599B2, EP3265817B1). P.P., G.DS.C., Z.T., P.R.B., and D.A.M. hold patents for the prediction of vaginal microbiota composition and inflammatory status (US2025/0027938A1). D.A.M. is a consultant for Calla Lily Clinical Care and Freya Biosciences. Other authors report no conflicts of interest.

## Abstract

Accurate and timely diagnosis of early pregnancy outcomes remains a clinical challenge, particularly in cases classified as pregnancy of unknown location, where ultrasound fails to confirm the location of the pregnancy. Delayed diagnosis can lead to serious complications, especially in ectopic pregnancies. This prospective cohort study aimed to characterise host-microbe interactions during early pregnancy in the vaginal mucosa by exploring the relationship between the bacterial community composition, immune response, and metabolic phenotype. We analyzed vaginal swabs from 91 women with pregnancy of unknown location, including 22 with viable intrauterine pregnancies and 69 with adverse outcomes. Metataxonomic profiling, immune assays, and direct on-swab metabolomics were performed. Adverse outcomes, especially ectopic pregnancies, were associated with reduced *Lactobacillus* abundance, increased bacterial diversity, mucosal inflammation, and elevated lipid species. Vaginal bleeding, a common symptom in PUL, significantly impacted immune and metabolic profiles and should be considered as a confounder in studies of the lower female reproductive tract. Multi-*omic* data integration highlighted metabolic phenotypes of immune response in the context of bacterial community composition, underscoring the influence of microbe-host interactions on mucosal health and pregnancy outcomes.

**IMPORTANCE:** Early detection of complications in pregnancy is essential for improving maternal health outcomes, yet current diagnostic tools often fail to identify ectopic or failing pregnancies in a timely manner. This study highlights how changes in the vaginal microbiome, inflammation, and metabolic signals are closely linked to adverse outcomes in early pregnancy. By showing that metabolic profiling can predict microbial and immune states even in the presence of vaginal bleeding, the research provides a foundation for developing rapid, non-invasive diagnostic tools. These insights could pave the way for more personalized and effective early pregnancy care.

## INTRODUCTION

Pregnancy of unknown location (PUL) is defined as a positive pregnancy test with no pregnancy seen on a transvaginal ultrasound. Subsequent pregnancy outcomes include intrauterine pregnancy (viable, VIUP or non-viable, NVIUP), failed PUL (FPUL), persisting PUL (PPUL), or ectopic pregnancy (EP)(1), with EP posing the highest risk of maternal mortality(2). The complex management of PUL has driven risk prediction models based on β-hCG and progesterone levels(3–5). The inclusion of additional clinical parameters such as vaginal bleeding, which is common in PUL, may improve model performance(6) however there remains the need for identification of novel biomarkers that reflect the underlying pathology of adverse early pregnancy outcome events.

We have previously reported that first-trimester miscarriage is associated with *Lactobacillus*-depleted vaginal microbiota and elevated levels of pro-inflammatory mediators(7)^-^(8). Increased prevalence of *Lactobacillus*-depleted vaginal microbiota has also been reported in women with PUL and tubal EP(9). Vaginal bacterial composition (VBC) is a major mediator of local immune and inflammatory responses(10–14) and the vaginal mucosal metabolome(13, 15–17), which are mechanistically implicated in the pathophysiology of adverse early pregnancy outcome such as miscarriage and EP(7, 18, 19). Characterization of reproductive tract microbiota-host interactions in adverse early pregnancy outcome therefore offers a strategy for identifying new biomarkers that may have clinical utility. However, in non-pregnant women, menstrual bleeding augments local immune responses(20, 21), and changes VBC(22, 23), and the vaginal mucosal metabolome(24, 25). The effect of vaginal bleeding during early pregnancy on these parameters is currently unknown.

In this study we use a multi-omic approach to characterize the vaginal microbiome, immune status, and metabolome in 91 women with PUL, to characterise mucosal microbe-host interactions in this population and determine whether molecular profiles can be used to predict subsequent pregnancy outcome.

## RESULTS

### Study participants

A total of 91 PUL cases were included in the study. This cohort consisted of 22 VIUP, 15 NVIUP, 26 FPUL, 20 EP, and 8 PPUL. VIUP cases served as the control group. No significant differences in key demographics such as age, gestation or ethnicity were observed between patient groups (Table 1, Supplementary Tables 1 and 2). Vaginal bleeding was reported and graded using Pictorial Blood Loss Assessment Chart (PBLAC) scores, with blood presence on swabs (PBOS) noted by clinicians at time of sample collection. There was strong agreement between PBLAC and PBOS (Pearson’s-χ2:p=8.83×10-7, Spearman’s:ρ=0.569, Fig. S1). Vaginal bleeding was associated with adverse early pregnancy outcomes, measured by both PBLAC (Pearson’s-χ2:p=3.80×10-4) and PBOS (Pearson’s-χ2:p=2.02×10-4, Table 1). Most VIUPs showed no bleeding (77%, PBLAC=0 and 95%, no PBOS), while bleeding was more common in adverse early pregnancy outcomes (76.7%, PBLAC score ≥1 and 46% PBOS), especially in FPUL cases (84%, PBLAC≥1 and 65% PBOS) (Supplementary Table 1 and Fig. S2).

**Table 1.** Demographics and clinical characteristics. Sub-comparison between viable and adverse PUL patient outcome groups. The adverse group consists of patient with non-viable intrauterine pregnancy (NVIUP), failed PUL (FPUL), ectopic pregnancy (EP) and persistent PUL (PPUL).

| Outcome | Viable, N = 22<sup>1</sup> | Adverse, N = 69<sup>1</sup> | p-value<sup>2</sup> |
| --- | --- | --- | --- |
| Maternal Age (years) | 31.0 (28.0, 34.5) | 35.0 (29.0, 39.0) | 0.080 |
| Ethnicity |  |  | >0.999 |
| Arab | 1 (4.5%) | 3 (4.3%) |  |
| Asian | 3 (14%) | 8 (12%) |  |
| Black | 2 (9.1%) | 5 (7.2%) |  |
| Caucasian | 15 (68%) | 47 (68%) |  |
| Other | 1 (4.5%) | 6 (8.7%) |  |
| Gestational age at presentation (days) | 32 (30, 35) | 40 (31, 47) | 0.027 |
| Unknown | 2 | 7 |  |
| PBLAC Vaginal bleeding |  |  | <0.001 |
| No bleeding (score 0) | 17 (77%) | 16 (23%) |  |
| Minimal bleeding (score 1) | 4 (18%) | 23 (33%) |  |
| Moderate bleeding (score 2) | 0 (0%) | 13 (19%) |  |
| Soaked sanitary towel (score 3) | 1 (4.5%) | 11 (16%) |  |
| Clots or flooding (score 4) | 0 (0%) | 6 (8.7%) |  |
| Presence of blood on swab |  |  | <0.001 |
| No | 21 (95%) | 37 (54%) |  |
| Yes | 1 (4.5%) | 32 (46%) |  |

<sup>1</sup>Median (IQR); n (%)
<sup>2</sup>Wilcoxon rank sum test; Fisher's exact test; Pearson's Chi-squared test

### Association of vaginal microbiota with pregnancy outcomes

Metataxonomic analysis of vaginal bacterial communities (VBC) detected 142 genera-level and 250 species-level taxa across the whole cohort. CSTs were assigned using the VALENCIA algorithm with 30.77% of patients being classified as CST I (*Lactobacillus crispatus*-dominated), 9.89% CST II (*Lactobacillus gasseri*-dominated), 36.26% CST III (*Lactobacillus iners*-dominated), 18.68% CST IV (mixed anaerobes), and 4.40% CST V (*Lactobacillus jensenii*-dominated). For genera level group classifications, CST IV was categorized as *’Lactobacillus*-depleted’ (LDEPL, 18.68%) and CST I, II, III, or V as *’Lactobacillus*-dominated’ (LDOM, 81.32%).

A higher proportion of VIUP had LDOM compared to women with adverse early pregnancy outcomes (Pearson’s-χ2:p=9.83×10-3, Fig. 1A, Fig. S3A), who were enriched for IV (Pearson’s-χ2:p=6.15×10-5, Fig. 1B, Fig. S3B) and higher Shannon α-diversity (Fig. S4A). Linear regression of clr-transformed species counts revealed higher relative abundance of anaerobic organisms (*Prevotella timonensis*, *Dialister micraerophilus*, *Streptococcus anginosus*, and *P. bivia*) in women with adverse early pregnancy outcomes (Fig. 1C) and lower *L. gasseri* levels (q<0.05, BH-FDR correction).

**FIG 1.**
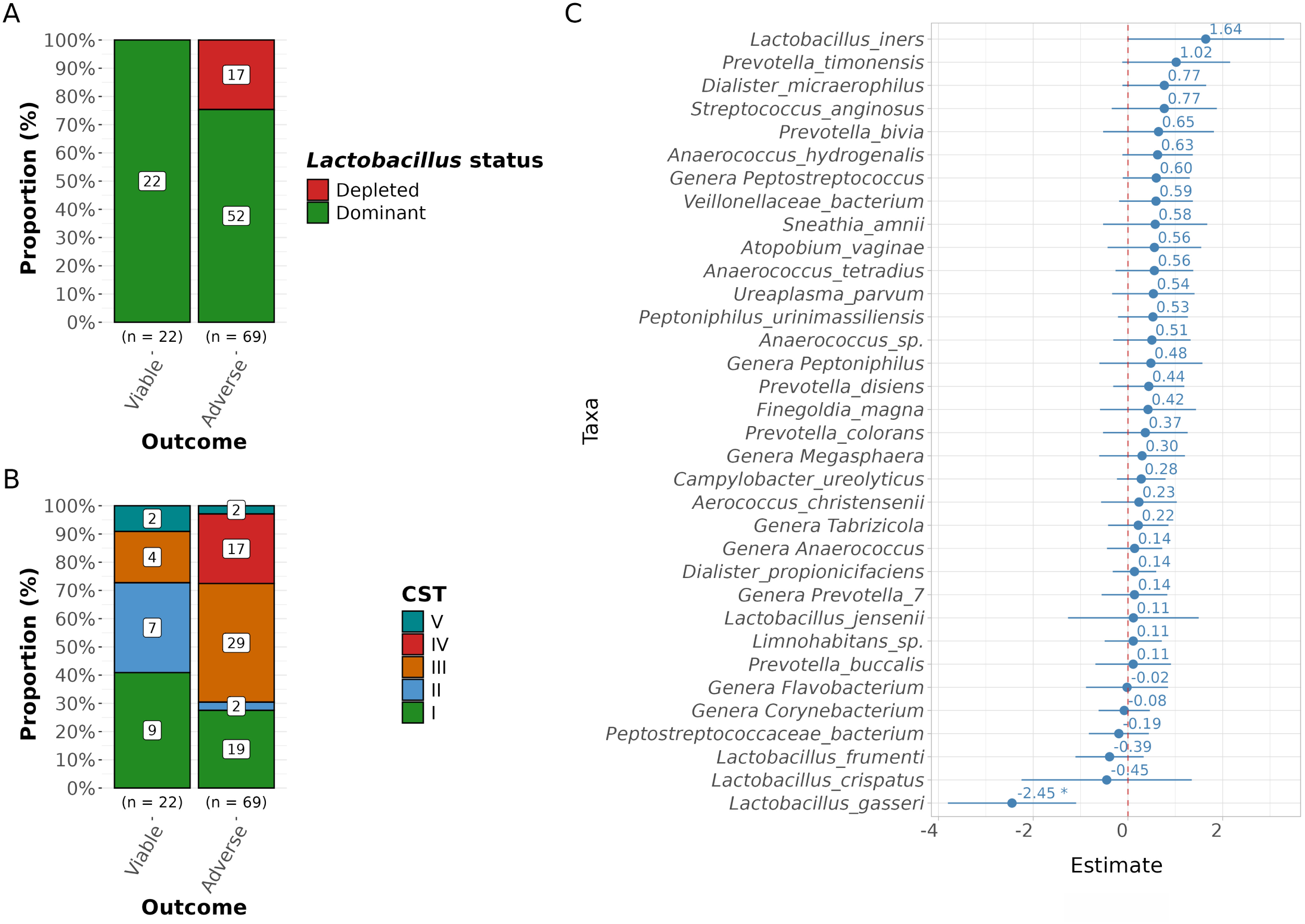
Association of vaginal bacterial communities with pregnancy outcome. Comparison between viable VIUP (n=22) and adverse early pregnancy outcome (n=69) outcome. Association of pregnancy outcome with **A)** prevalence of *Lactobacillus* dominance/depletion (*red: L. depleted; green: L. dominant)* **B)** community state types (*CST I: green, II: light blue; III (orange), IV (red), V dark blue)* and **C)** individual bacterial species. BH-FDR q<0.05 denoted by *.

In sub-analyses, EP was shown to have higher LDEPL prevalence (Pearson’s-χ2:p=5.52×10-3, Fig. S5A), enrichment for CSTs III and IV (Pearson’s-χ2:p=2.86×10-4, Fig. S5B), and higher Shannon α-diversity (Fig. S4B) compared to VIUP. At the species level, *L. iners*, *Sneathia amnii* and *P. timonensis* showed strongest association with EP (Fig. S5C). To control for bleeding, we analyzed PBLAC and PBOS variables and found a balanced distribution of vaginal bleeding across CST groups (Fig. S6A and S6B), except for samples with PBLAC=4, which were enriched for CST IV with higher Shannon α-diversity (Fig. S6C and S7C). Linear regression did not identify strong bacterial signatures of vaginal bleeding after BH-FDR correction (Fig. S8).

### Immune signatures linked to pregnancy outcome and vaginal bleeding

To evaluate the relationship between immune mediator concentrations, pregnancy outcomes, and vaginal bleeding, a PCA model was applied to log-transformed immune mediator concentrations. The first two principal components accounted for 52.27% of total variance, which was explained by elevated levels of TNF-α, GM-CSF, MMP-1, CCL2/MCP-1, IL-6, and LIF, and were associated with adverse early pregnancy outcomes (Fig. S9A, PERMANOVA:p=1.00×10-3). However, this association was confounded by vaginal bleeding (PBLAC≥1, PERMANOVA:p=3.40×10-3; PBOS, PERMANOVA:p=2.00×10-4), and adjusting for bleeding nullified the significance (Fig. 2). Linear regression also identified an immune signature associated with adverse early pregnancy outcomes, characterized by increased levels of 10/15 immune markers (q<0.05, Fig. 2B). However, these differences were no longer significant after adjusting for bleeding (Fig. 2C, Fig. S9B and S9C). Further analysis identified significant positive correlations (q<0.05) between immune markers and bacteria associated with CST IV (Fig. 3). For example, *P. bivia* correlated with TNF-α (r²=0.19) and MMP-1 (r²=0.1), *Sn. amnii* with IL-1α (r²=0.13) and LIF (r²=0.13), and *S. anginosus* with TNF-α (r²=0.2), IFN-γ (r²=0.12), IL-10 (r²=0.14), ICAM-1 (r²=0.096), and MMP-1 (r²=0.11). In contrast, *L. crispatus*, *L. gasseri*, and *L. jensenii* showed inverse correlations with most immune markers, but these were not significant after BH-FDR correction (q<0.05).

**FIG 2.**
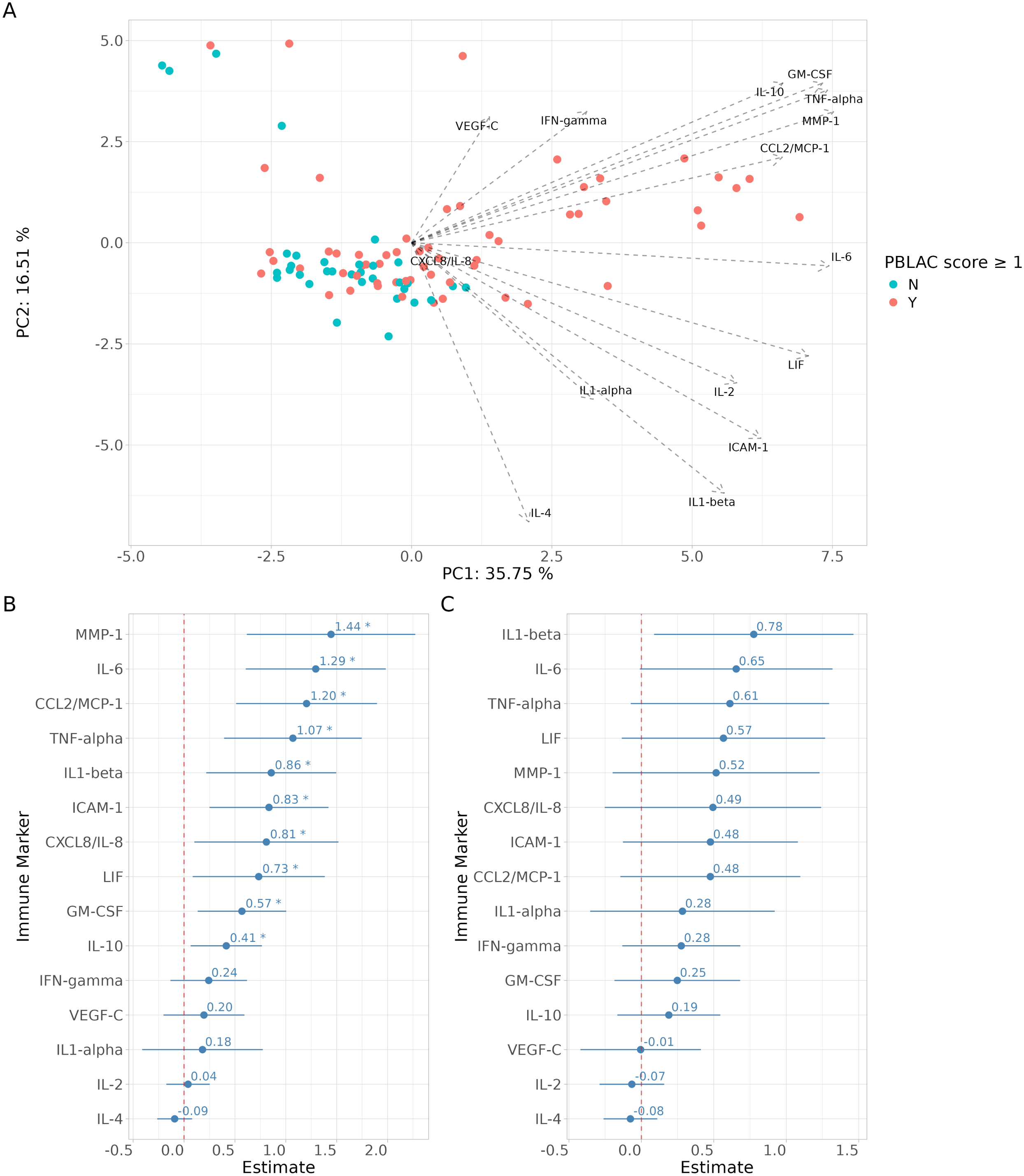
Association of immune signatures with vaginal bleeding and pregnancy outcome. **A)** Unsupervised PCA analysis of the log-transformed immune marker concentrations. Sampling points were colored by vaginal bleeding state (blue: No bleeding; red: PBLAC≥ 1). Immune markers associated with adverse early pregnancy outcome **B)** without confounder adjustment and **C)** with adjustment for bleeding using PBOS (*BH-FDR q<0.05).

**FIG 3.**
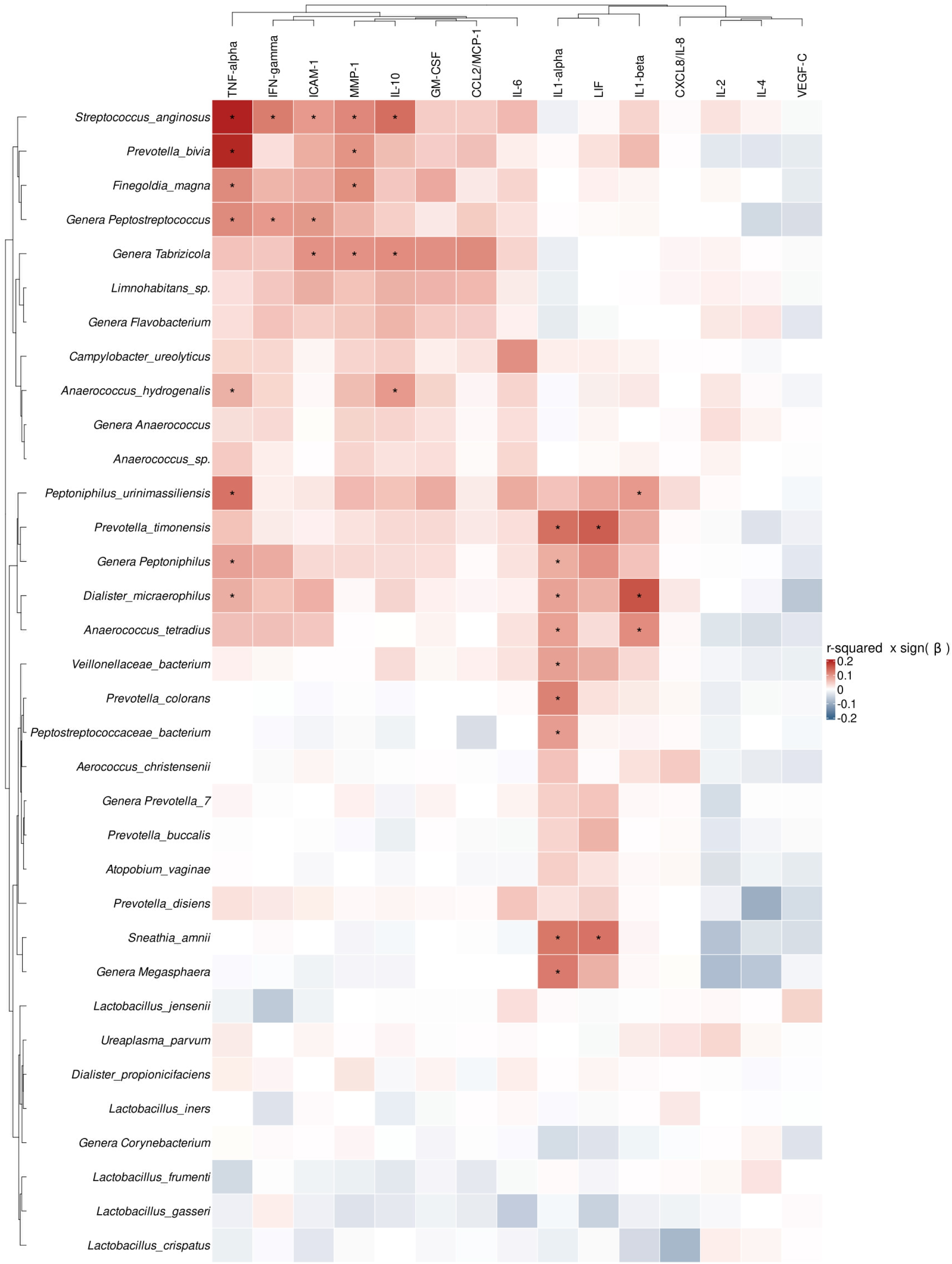
Associations between individual bacterial taxa and immune signatures. Ward hierarchical clustering heatmap plot representing correlations between immune markers and bacteria taxa. Cells are coloured by the variance explained in cytokine levels (r^2^) multiplied by the sign of the regression coefficient (β*) (red: max=0.2; blue: max=-0.2*, *BH-FDR, q<0.05).

### Vaginal metabolites correlate with VBC but not pregnancy outcomes

To investigate the relationship between vaginal metabolites and VBC, a PCA model with 5 components (explaining 50.22% of the variance) was fitted to DESI-MS metabolite profiles and the score plots coloured according to CST (Fig. 4A) or PBOS (Fig. 4B). PERMANOVA identified VBC as the main source of variability (R²=0.116, p=1.10×10⁻³), followed by vaginal bleeding (R²=0.0543, p=1.40×10⁻³ for PBOS, not significant for PBLAC≥1). No significant association with pregnancy outcome was observed. Multivariate random forest models built using metabolomic data were able to predict VBC, bleeding, and outcome. Models could robustly discriminate VBC at genera-level (LDOM vs. LDEPL, 7-fold CV AUC=0.84, Fig. 4C) and between CSTs I, III, and IV (Fig. S10). Vaginal bleeding prediction was robust using the PBOS variable (7-fold CV AUC=0.77, Fig. 4D, Fig. S11). Linear regression identified metabolic features associated with VBC (LDOM vs. LDEPL) and vaginal bleeding (Fig. S12). Metabolite biomarkers increased in LDEPL samples included fatty acids, monoradylglycerols (MG), and diradylglycerols (DG), while LDOM samples had increased small molecules like citrate, glutamate, galactose, and 5-formyluridine (Fig. 5A). Bleeding was characterized by increased lipids (PS, PC, DG, CER, SM) (Fig. 5B). Analyses distinguishing VIUP from adverse early pregnancy outcome did not yield robust predictive models or significant metabolic markers after BH-FDR correction. Hierarchical clustering of 13 cytokine mediators revealed two groups (Fig.6). Group 1 (IL-6, MMP-1, CCL2/MCP-1, TNF-α, ICAM-1, GM-CSF, IL-10) was associated with lipid classes (PC, PE, SM, CER, PI) and correlated with PBOS. Group 2 (IL-1β, CXCL-8/IL-8, IL-1α, LIF, IL-2, IFN-ɣ) was linked to oxypurinol, CER(36:2), and Galabiosylceramide(d34:1), with the latter also associated with VBC. Metabolites negatively associated with increased cytokine expression included small molecules, fatty acids, lyso-lipids, and monoglycerides, most of which positively correlated with VBC.

**FIG 4.**
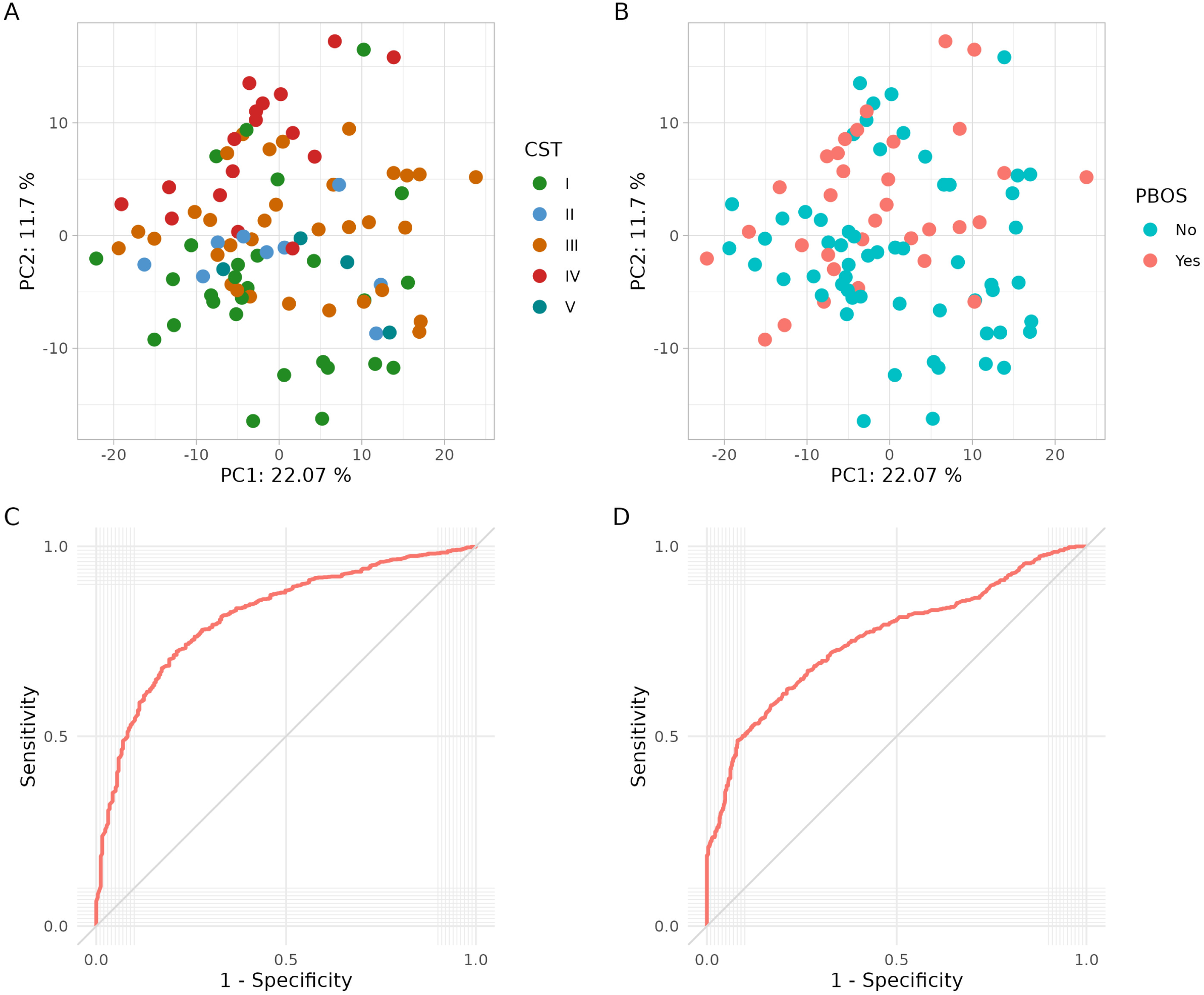
Predictive capacity of VBC and PBOS using vaginal metabolites. Univariate PCA analysis of vaginal metabolites. PCA score plot of the first 3 principal components shows good separation of samples on the basis of **A)** CST groups (I: green, II: light blue, III: orange, IV: red, V: dark blue) and to a lesser extent, **B)** blood present on swab (red: presence, blue: absence). Random Forest classifier analysis and ROC curve analyses indicate vaginal metabolites offer good classification performance for **C)** *Lactobacillus* dominant vs depleted status (AUC=0.84) and **D)** PBOS (AUC=0.77).

**FIG 5.**
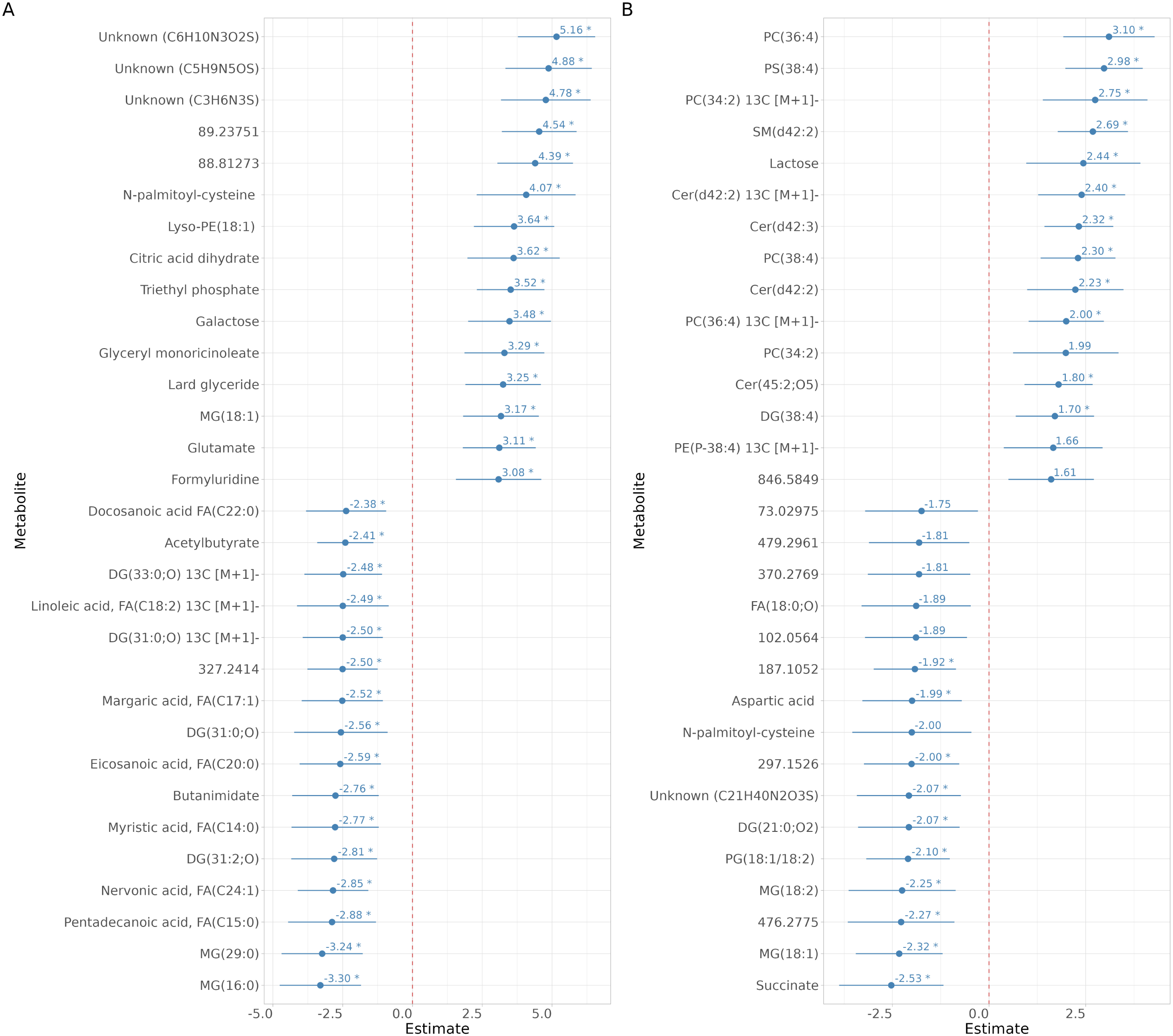
Vaginal metabolite biomarkers associated with vaginal bacterial composition and blood presence on swabs. Top 30 statistically discriminative vaginal metabolite markers associated with **A)** *Lactobacillus* dominant vs depleted status and **B,** with blood presence on swabs. *BH-FDR, q<0.05.

**FIG 6.**
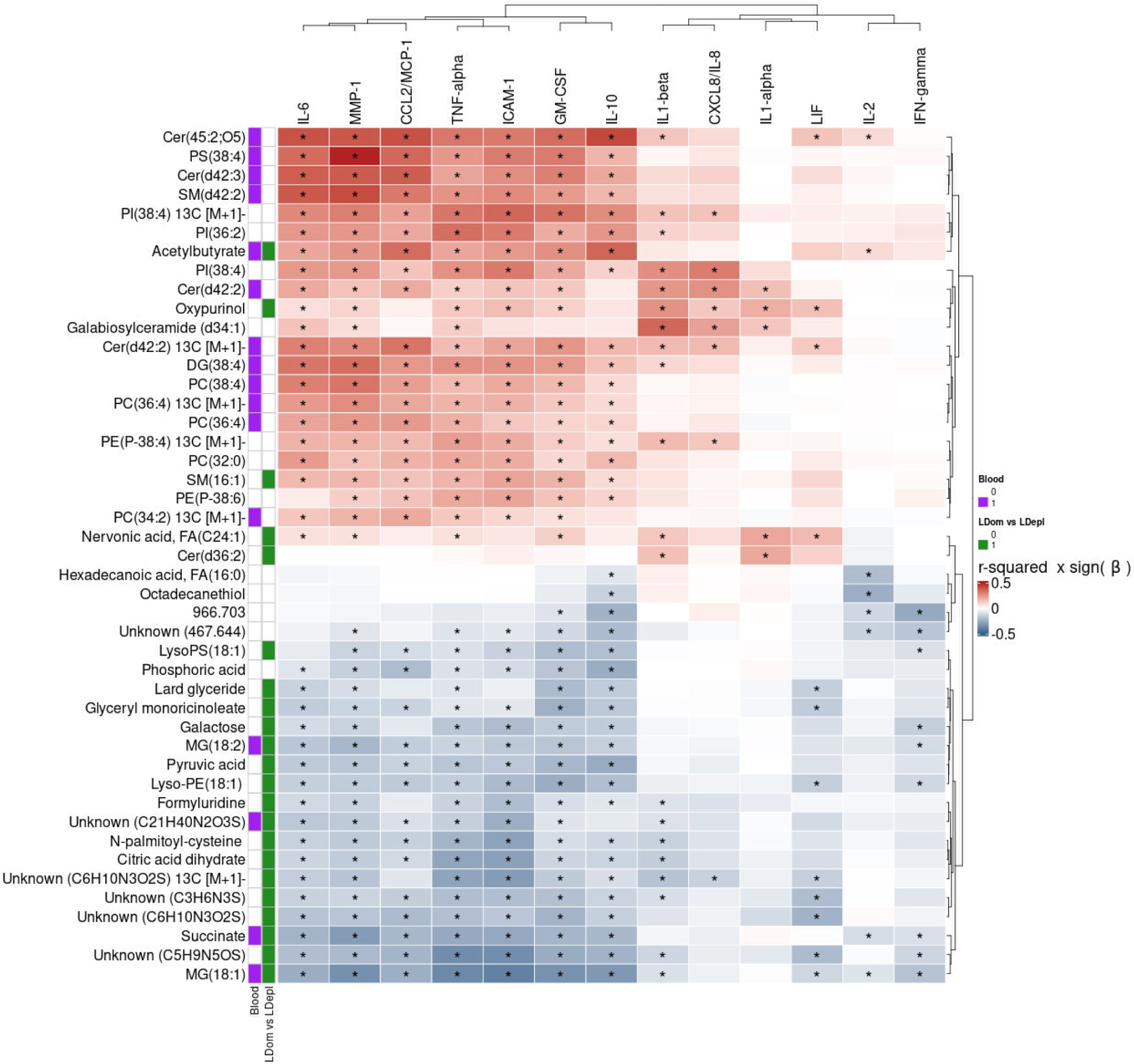
Vaginal metabolite biomarkers associated with immune signatures and VBC. Ward hierarchical clustering of vaginal immune markers and metabolite features. Correlation values of metabolites were coloured and range between 0.5 (red) and -0.5 (blue). Metabolites exhibiting co-correlations with blood (purple) and *Lactobacillus* dominant vs depleted status (green) are also highlighted (*BH-FDR, q<0.05).

### Integration of vaginal microbiota, immune and metabolic profiles

The three datasets were analyzed using the MOFA+ model(26) to investigate shared variance among bacterial composition, immune response, metabolic profiles, pregnancy outcome, and vaginal bleeding (Fig. 7). A MOFA+ model with 5 factors accounted for 16.29%, 26.80%, and 48.92% of the variance in metataxonomic (16S), immune, and metabolomes, respectively (Fig. 7A). Factor 1 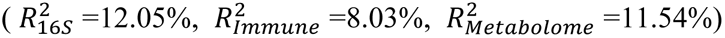 correlated with PUL outcome (r²=0.12) and CST (r²=0.41). Factor 3 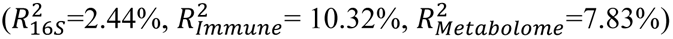 correlated with bleeding (PBLAC ≥1 r²=0.069, blood visible on swab r²=0.18) and outcome (r²=0.124) (Fig. 7B). Factor 1 was driven by the LDEPL state, increased pro-inflammatory markers (TNF-α, IL-1β, ICAM-1, IL-1α, LIF, MMP-1, IL-6), and metabolic features associated with LDEPL (e.g., fatty acids like FA(C20:0) and FA(C17:0)) (Fig. 7C). Factor 3 showed an increase in most immune markers, particularly CCL2/MCP-1, MMP-1, TNF-α, and a higher abundance of glycerophosphocholines (e.g., PC(36:4) and PC(36:3)) (Fig. 7D).

**FIG 7:**
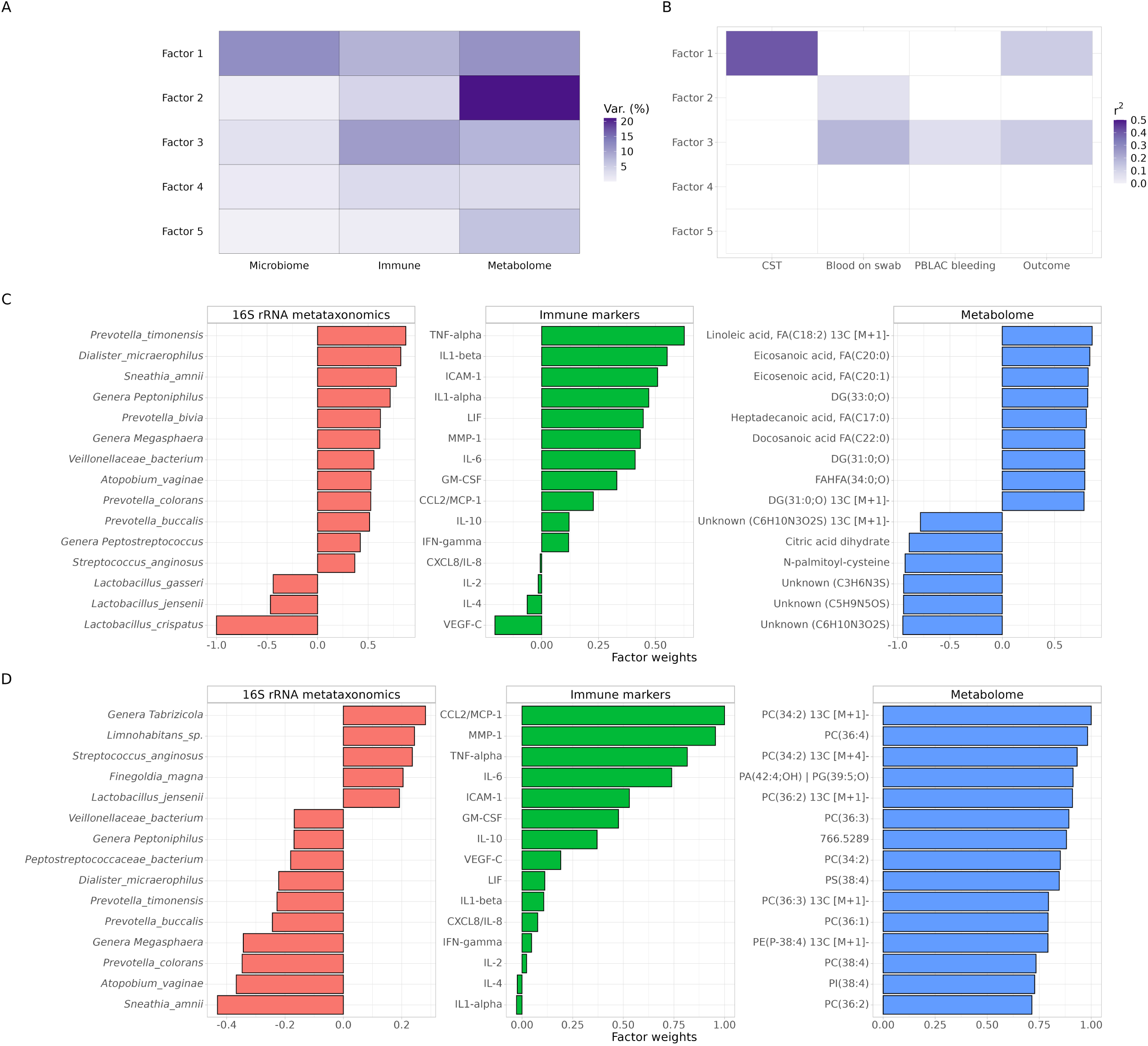
Multi-omic factorial analysis of most significant covariate factors. Assessment of the proportion of variance explained by each of the top 5 factor between **A)** microbiome, immune and metabolome and between **B)** CST, PBOS, PBLAC bleeding and outcome. Top 15 absolute factor weights per data block for **C)** Factor 1 and **D)** Factor 3 for 16S rRNA meta-taxonomic (left), immune markers (centre) and metabolomics (right) data.

## DISCUSSION

This study presents new insights into the complex interplay between VBC, host immune responses, and metabolic activity in early pregnancy, focusing on women with PUL. By integrating metataxonomic, immune, and metabolic profiling, we identified distinct biological signatures associated with adverse early pregnancy outcome particularly EP and highlight the potential of vaginal metabolomics as a tool for early risk stratification.

Our results show that adverse early pregnancy outcomes are associated with significant alterations in the vaginal microbiome. Specifically, we observed a consistent depletion of *Lactobacillus* species and an increase in microbial diversity in women who subsequently experienced miscarriage or EP. This is consistent with previous findings that link *Lactobacillus*-dominant vaginal ecosystems with reproductive tract health and successful pregnancy. *Lactobacillus* species, particularly *L. crispatus* contribute to a protective mucosal environment through the production of lactic acid and antimicrobial compounds(27), maintaining low pH(28) and suppressing pro-inflammatory cytokines such as IL-6, IL-8, and TNF-α(29). While the precise mechanism by which vaginal microbiota-host interactions shape the risk of adverse early pregnancy outcome remains to be fully elucidated, it is possible that a VBC depleted in *Lactobacillus* and increased in bacterial diversity leads to disruption of the highly regulated immune and inflammatory events involved in endometrial implantation, predisposing a woman to developing an EP or other adverse pregnancy outcomes(30). Recent evidence suggests the translocation of vaginal bacteria and their metabolites to the endometrial environment can activate inflammatory pathways associated with pregnancy complications(31). Consistent with this, we observed an enrichment of *Sneathia* species in women with EP, which is known to cause inflammation within the female genital tract(32–34). Alternatively, aberrant implantation during EP may lead to other bleeding events or changes in hormonal levels that subsequently lead to alterations in VBC.

Indeed, one of the major confounding factors we identified was the presence of vaginal bleeding. Bleeding is common in PUL cases and strongly influenced both immune and metabolic readouts from vaginal swabs. Immune mediator concentrations initially appeared to be associated with pregnancy outcome, but these associations were no longer significant after adjusting for the presence of blood. Similarly, bleeding influenced the metabolic profile of cervicovaginal fluid, leading to increased abundance of certain lipid species known to derive from blood components. These findings emphasize the importance of accounting for bleeding when analyzing vaginal samples for diagnostic or research purposes.

Despite the confounding effect of bleeding on immune and metabolic markers, our analysis demonstrated that direct metabolic profiling of vaginal swabs using desorption electrospray ionisation-mass spectrometry (DESI-MS) could robustly identify bacterial states and reflect immune activation, independent of blood contamination. Metabolic features, particularly lipid-related species, showed strong correlations with bacterial composition and mucosal inflammation. Multi-omic factor analysis identified two principal biological axes: one capturing host immune and metabolic responses to *Lactobacillus* depletion and another reflecting immune-metabolic changes associated with bleeding. Interestingly, neither axis alone was strongly predictive of pregnancy outcome, underscoring the complexity of these interactions and the likely involvement of additional host and microbial factors.

These findings have important implications for both clinical practice and research. From a clinical perspective, the ability to rapidly assess vaginal microbial and immune-metabolic states using minimally invasive swab-based methods could enhance early diagnosis and risk stratification in early pregnancy. Current diagnostic approaches rely heavily on serial ultrasound and serum human chorionic gonadotropin (hCG) measurements, which may delay diagnosis and appropriate intervention. The identification of vaginal microbial signatures that precede clinical outcome offers an opportunity to detect high-risk pregnancies earlier, potentially improving outcomes through closer monitoring or timely treatment. Given that all samples in this study were collected prior to definitive clinical diagnosis, our data support the potential of microbial and metabolic biomarkers as early predictors of adverse early pregnancy outcomes, particularly EP.

From a research standpoint, this study adds to the growing body of evidence linking vaginal dysbiosis to poor reproductive outcomes. However, the underlying mechanisms remain poorly understood. It is still unclear whether these microbial shifts directly contribute to adverse outcomes, or whether they arise in response to other physiological disruptions. Possible mechanisms include translocation of bacteria or microbial products to the endometrium or fallopian tubes, modulation of local immune responses, or systemic inflammatory signalling affecting implantation and embryo development. These hypotheses remain difficult to test due to the absence of reliable animal models and practical and ethical limitations in studying human implantation. Further mechanistic studies, including those incorporating longitudinal sampling from pre-conception through early gestation, will be necessary to clarify causality.

Strengths of this study include the analysis of a well-characterised cohort with detailed information on vaginal bleeding. The use of 3 complementary molecular profiling techniques provides a comprehensive view of bacteria and host interactions in PUL. The use of metataxonomics enabled the identification and evaluation of a wide range of commensal and pathogenic vaginal bacteria. However, the single 38F forward primer used amplification of the V1-V2 region has poor detection of *Gardnerella vaginalis*, which has been reported to be significantly elevated in other studies of EP(35). Other limitations include the relatively low number of samples and the absence of pre-conception samples, which would help clarify whether *Lactobacillus* depletion precedes or results from adverse early pregnancy outcome and determine if high levels of MMP-1 and other pro-inflammatory markers are due to pathogenesis or blood contamination. The mechanisms leading to the development of an EP are poorly understood but *Chlamydia trachomatis* has been repeatedly implicated(36). We lacked data on current *C. trachomatis* infection, which is key to determining if the bacterial profile is independent of pelvic chlamydial infection(37).

In summary, our findings indicate that adverse early pregnancy outcomes are associated with a shift in the vaginal microbiome towards a *Lactobacillus*-depleted, high-diversity state, accompanied by increased mucosal inflammation and altered metabolic profiles. While vaginal bleeding complicates the interpretation of immune data, metabolic profiling via DESI-MS provides a robust readout of bacterial community state and host response. These insights support the potential use of vaginal metabolic profiling as a rapid, non-invasive diagnostic approach for early pregnancy complications. Future studies should focus on validating these findings in larger, diverse cohorts and elucidating the causal pathways linking microbial dysbiosis to early pregnancy loss.

## MATERIALS AND METHODS

### Study population

Eligible women presenting to the Early Pregnancy Assessment Unit at Queen Charlotte’s and Chelsea Hospital (London, UK), classified with PUL based on initial transvaginal ultrasound scan, were invited to participate. Exclusion criteria included age <18 or >50 years and positive HIV or hepatitis B/C status. Detailed medical and gynecological histories were obtained. Final pregnancy outcomes were determined at 11-14 weeks gestation. Each participant provided two mid-vaginal swabs collected using a BBLTM CultureSwabTM with liquid Amies (Becton Dickinson) and a Transwab MW170 (Medical Wire & Equipment) at a single time point. The swab placed in culture medium was used for microbiome analysis, while the dry swab was reserved for initial, non-destructive surface direct metabolomic profiling, followed by cytokine analysis. Severity of bleeding was assessed using the Pictorial Blood Loss Assessment Chart (PBLAC)(38, 39). Swabs were immediately chilled on ice and stored at −80°C.

## 16S rRNA gene amplicon sequencing

Bacterial DNA was extracted from BBL CultureSwab swabs using a QiAmp Mini DNA kit (Qiagen) as previously described^19^. The V1-V2 hypervariable regions of 16S rRNA genes were PCR-amplified using 28F-GAGTTTGATCNTGGCTCAG and 388R-TGCTGCCTCCCGTAGGAGT primers. Sequencing was conducted by RTL Genomics (Lubbock, TX) and primer sequences were trimmed using cutadapt(40) (v2.8). Trimmed paired reads were processed in QIIME2(41) with the DADA2(42) denoising algorithm to obtain amplicon sequence variant (ASV). A custom naive Bayes classifier(41) trained on the Silva SSU (version 138)(43) reference database was used to assign taxonomy to the ASV sequences.

Diversity metrics were calculated with QIIME2 diversity plugin. Classification of samples based on their vaginal microbiota community state types (CST) was performed with the VALENCIA(44) nearest-centroids classifier.

### Direct-swab DESI-MS metabolite profiling

The method and instrument setting parameters used for the direct analysis of vaginal swabs by desorption electrospray ionization mass spectrometry (DESI-MS) are detailed elsewhere(45). Briefly, a DESI-MS source was used with a LTQ Orbitrap Discovery mass spectrometer (Thermo Scientific). The vaginal mucous was absorbed from a small area of the swab tip surface by gently desorbing the analytes with charged droplets of methanol/water (95:5, v/v) mixture with a solvent flow rate of 10μl per minute. For each sample, 30 scan mass spectra (*m/z* 50–1000, R=30,000 (FWHM)) were recorded in the negative ion mode. RAW files were converted to .mzML format using proteoWizard^45^. The mass spectrum data underwent processing with maldiQUANT(46). Target *m/z* features were tentatively identified through searches in HMDB and Lipidmaps with a 5ppm search tolerance. Structural elucidation was further conducted via MS/MS experiments. Swab analysis per sample was brief, lasting approximately 2–5 minutes. After analysis, the samples were returned to the freezer for storage until further processing for immune profiling.

#### Immunoprotein-profiling of vaginal swabs

Swabs were thawed on ice and re-suspended in 350ml of PBS solution containing protease inhibitor (5μL/ml; Sigma-Aldrich) and vortexed for 1 minute. The swab was inverted and transferred into a new microcentrifuge tube before being centrifuged at 8000xg for 10 minutes. Both suspensions were pooled before being centrifuged at 13000x*g* for 10 minutes. Quantification of immune mediators was performed using the Human Magnetic Beads Luminex Screening Assay 15-kit (Luminex Corporation)(IL-1α, IL-1β, IL-2, IL-4, IL-6, IL-8 (CXCL-8), IL-10, GM-CSF, MCP-1 (CCL2), TNF-α, IFN-γ, VEGF-C, ICAM-1, LIF and MMP-1) with a Bio-Plex 200 system (Bio-Rad Laboratories).

### Statistical analysis

Baseline characteristics were compared using Wilcoxon’s rank-sum test for continuous variables and Fisher’s exact test for categorical variables. Raw counts from metataxonomics data were clr-transformed. Immune marker concentrations and MS feature intensities were log-transformed after adding a constant offset (c=1). Linear models were fitted using base R *lm* method, with r2 measures, Welch’s t-test, and ANOVA test. Regression analyses modeled each variable as a function of outcome and/or bleeding score. Associations between bacterial taxa and immune species were calculated from linear models with log-transformed immune marker concentrations as dependent variables and clr-transformed counts as independent variables. Immune-metabolic associations regressed log-transformed immune marker concentrations against log-transformed MS feature intensities. The Benjamini-Hochberg method corrected for false discovery rate (BH-FDR) in all linear model analyses. When correlating bacterial taxa or metabolites with cytokines, BH-FDR correction was applied to each immune mediator separately. PCA models were fitted from clr-transformed counts or log-transformed variables. PERMANOVA models and marginal tests used Euclidean distance and 5000 permutations. Multivariate random forest classifiers were trained and cross-validated with 15 repeats of 3-fold stratified CV, increasing to 7 folds for bleeding status and clinical outcomes. All models used 1000 decision trees and an mtry value of 1/3 of the number of variables. ROC curves were calculated, and predictive RF classifiers were validated using 1000 permutation tests. For the multi-omics factor analysis (MOFA+) model, 5 factors were selected to decompose the three data blocks, all modeled with Gaussian noise.

## DATA AVAILABILITY

Public access to sequence data sets generated in this study along with accompanying metadata can be obtained from the Sequence Read Archive of the European Nucleotide Archive (PRJEB72306), the MetaboLights metabolomic repository (MTBLS10098) and Github (https://github.com/Gscorreia89/abpep-desi-pul).

## ETHICS APPROVAL

This study was approved by the National Health Service (NHS) National Research Ethics Service (NRES) North of Scotland Research Ethics Service (REC 14/NS/1078).

## ACKNOWLEDGEMENTS

Financial support for this study was provided by the Tommy’s National Centre for Miscarriage Research (P62774) and the UK National Institute for Health Research Biomedical Research Centre (P45272). G.DS.C., J.R.M, Z.T., P.R.B. and D.A.M. are supported by the March of Dimes European Prematurity Research Centre at Imperial College London. S.B. was supported by NIHR CLAHRC North West London (RDIP033). The Division of Digestive Diseases and receives financial support from the National Institute of Health Research (NIHR) Imperial Biomedical Research Centre (BRC) based at Imperial College London and Imperial College Healthcare NHS Trust.

